# Soil microbial diversity impacts plant microbiota more than herbivory

**DOI:** 10.1101/2020.09.30.320317

**Authors:** Antonino Malacrinò, Alison J. Karley, Leonardo Schena, Alison E. Bennett

## Abstract

Interactions between plants and microbiomes play a key role in ecosystem functioning and are of broad interest due to their influence on nutrient cycling and plant protection. However, we do not yet have a complete understanding of how plant microbiomes are assembled. Here, we tested and quantified the effect of different factors driving the diversity and composition of plant-associated microbial communities. We manipulated soil microbial diversity (high or low diversity), plant species (*Solanum tuberosum* or *Solanum vernei*), and herbivory (presence or absence of a phloem-feeding insect *Macrosiphum euphorbiae*), and found that soil microbial diversity influenced the herbivore-associated microbiome composition, but also plant species and herbivory influenced the soil microbiome composition. We quantified the relative strength of these effects and demonstrated that the initial soil microbiome diversity explained the most variation in plant- and herbivore-associated microbial communities. Our findings strongly suggest that soil microbial community diversity is a driver of the composition of multiple associated microbiomes (plant and insect), and this has implications for the importance of management of soil microbiomes in multiple systems.

## Introduction

Microbiomes can be considered an extension of the plant genome (Berg *et al.*, 2014; Schlaeppi and Bulgarelli, 2015; Rosenberg and Zilber-Rosenberg, 2016; Levy *et al.*, 2018). While their functional importance has been widely dissected in the last two decades of microbiome research, how plant microbial communities assemble, respond to environmental stimuli, and interact with their host remains to be determined (Cordovez *et al.*, 2019; Saikkonen *et al.*, 2020). In addition, research has rarely examined or compared multiple drivers and, to the best of our knowledge, no study has tested the relative strength of different drivers of microbial community composition.

There have been a number of studies identifying individual factors that drive the microbiome composition of plants and their associated organisms and environments. It is well established that plant microbiota structures mainly by plant compartment (e.g. different plant organs, and diverse between endosphere and ectosphere) (Trivedi *et al.*, 2020). Plant genotype and developmental stage have been shown to influence the composition of both plant and soil microbiomes (Wagner *et al.*, 2016). Soil microbiome composition has also been shown to shape plant microbiome composition (Cordovez *et al.*, 2019), and plant pathogens and herbivory produce compositional shifts in plant-associated microbial communities (Lareen *et al.*, 2016). However, we still know little about the relative strength of these factors in shaping plant microbiomes.

Soil provides microbial inoculum and sets the conditions for both plant and microbial growth (Schlaeppi *et al.*, 2014). While different plant tissues can develop distinct microbiomes, soil provides an important reservoir of microbial inoculum for both the phyllosphere and rhizosphere (Bai *et al.*, 2015). The overlap of soil and plant microbiota has been found in a variety of plants, for example *Saccharum officinarum* (de Souza *et al.*, 2016), *Boechera stricta* (Wagner *et al.*, 2016), *Vitis vinifera* (Zarraonaindia *et al.*, 2015; Mezzasalma *et al.*, 2018) and in the biofuel crops *Panicum virgatum* and *Miscanthus x giganteus* (Grady *et al.*, 2019). This suggests that soil microbial communities can represent a major factor shaping plant microbiomes in both above and belowground plant compartments. Furthermore, it has been shown that soil microbial community can influence the feeding behaviour of insect herbivores (Badri *et al.*, 2013). However, only one study to our knowledge has reported that soil microbial community can directly influence both the aboveground microbiota of plants (*Taraxacum officinale*) and the microbiota of an insect herbivore (*Mamestra brassicae*), showing overlap between the microbial communities of the insect and soil (Hannula *et al.*, 2019). However, it was unclear whether the influence of soil microbiome on the caterpillar’s microbiome was due to passive transfer (e.g. microbe dispersal when watering) or an active colonisation mechanism (Hannula *et al.*, 2019). Thus, there is potential for the influence of soil microbial communities on plant-associated microbial communities to extend beyond their plant host.

Plant species and genotype also contribute to the composition of multiple plant-associated microbiota (Turner *et al.*, 2013). Plant microbial communities assemble differently in different organs, and their composition varies according to plant phylogeny (Dastogeer *et al.*, 2020). Plant traits (tissue morphology and physicochemical properties) and resources might drive these plant-genotype effects on plant microbiota (Fitzpatrick *et al.*, 2018; Dastogeer *et al.*, 2020). For example, an analysis of 30 species of angiosperms revealed differences in the diversity and composition of root microbiomes across plant species (Fitzpatrick *et al.*, 2018). Similarly, the rhizosphere microbiota of wheat, maize, tomato and cucumber each had unique microbial communities (Ofek *et al.*, 2014), and leaf and root microbial communities of *Agave* species clustered according to the host plant species (Coleman-Derr *et al.*, 2016). Furthermore, several studies reported that host plant identity is an important factor in the assembly of insect herbivore-associated microbial communities (Colman *et al.*, 2012; Malacrinò, 2018). For example, host plant species influenced the composition of the microbiome associated with the herbivores *Ceratitis capitata* (Malacrinò *et al.*, 2018) and *Thaumetopoea pytiocampa* (Strano *et al.*, 2018). Therefore, plant species identity has an effect on the communities of microorganisms in the rhizosphere, living in the different plant organs, and even within plant herbivores.

Only a few studies have tested the effects of herbivory on plant microbiomes. For example, whitefly infestation of pepper plants led to an increased proportion of Gram-positive bacteria in the rhizosphere (Yang *et al.*, 2011), and aphid herbivory on pepper plants increased the abundance of *Bacillus subtilis* and decreased that of the pathogen *Ralstonia solanacearum* in roots (Lee *et al.*, 2012). While whitefly herbivory shifted the rhizosphere microbiome composition in pepper plants (Kong *et al.*, 2016), there was no effect of aphid herbivory on the rhizosphere microbiota of Brassica oleracea var. *capitata* (O’Brien *et al.*, 2018). Herbivore attack may alter plant root exudates which, in turn, promote the assembly of beneficial rhizosphere microbiota (Hu *et al.*, 2018; Rolfe *et al.*, 2019). Thus, herbivory can alter plant and rhizosphere microbiomes, but the relative impact of herbivory versus plant species or initial soil diversity on plant-associated microbiomes has not been investigated.

To date most studies have focused on one or two drivers of microbiome composition (e.g., soil microbiota, plant species or herbivory) and, to the best of our knowledge, no study has tested the comparative strength of multiple biotic drivers expected to shape above- and belowground microbiomes *in vivo*. This represents a major gap in our understanding of the relative importance of factors determining microbiome assembly. Here we ask how plant microbiome composition is shaped by three different major drivers of plant-associated microbial communities — soil microbial diversity, plant species, and herbivory — and whether they have equal impact on plant-associated microbiomes. By manipulating insect herbivory (presence/absence), plant species identity and soil microbial diversity in a microcosm system, and by quantifying their effects on both bacterial and fungal plant-associated microbiome composition, we tested the relative strength of these three biotic factors in shaping rhizosphere, plant (root and shoot), and herbivore microbiomes. As outlined above, there is evidence of overlaps between soil and plant microbiota, thus we hypothesize that soil microbial diversity is the major driver structuring plant microbiomes at different compartments. Previous studies reported that, within the same plant species, geographical location greatly contributes to assemble the plant microbiota (Christian *et al.*, 2016; Lin *et al.*, 2020). Similarly, there is evidence that soil microbial community can influence herbivores (Badri *et al.*, 2013) and their microbiota (Hannula *et al.*, 2019). Thus, we hypothesize that plant species and herbivory will impact plant-associated microbial communities, but their magnitude would be lower than soil microbial diversity.

## Methods

### Experimental design

In this study, we used a 2 × 2 × 3 factorial design to test our hypothesis. We grew two *Solanum* species (*Solanum tuberosum* and *Solanum vernei*) in soil with different microbial diversities: high diversity and low diversity (see below). To evaluate the effects of herbivory on plant and rhizosphere microbiota, we infested plants (within each ‘soil’ × ‘plant species’ combination) with two clonal lines of the polyphagous aphid species *Macrosiphum euphorbiae* (potato aphid); uninfested plants served as a control. Each treatment combination of plant species (*n*=2), soil microbial diversity (*n*=2), and aphid clonal line and presence/absence (*n*=3) was replicated five times, involving 60 plants in total.

### Study System

*Solanum tuberosum* (genotype TBR-5642) and *Solanum vernei* (genotype VRN-7630) seeds were obtained from the Commonwealth Potato Collection at The James Hutton Institute (Dundee, Scotland, UK). Seeds were germinated in steam-sterilized coir, and then transplanted to the experimental pots after 3 weeks.

We used two aphid clones of *M. euphorbiae* (AK13/08 and AK13/18) previously collected in the field (James Hutton Institute, Dundee, UK — 56.457 N, 3.065 W) and reared for several generations on excised leaves of *Solanum tuberosum* (cv. Desirée) in ventilated cups at 20 °C and 16:8 h (light:dark).

All inoculum was prepared from soil collected from an uncultivated field at the James Hutton Institute (56.457 N, 3.065 W) (Bennett *et al.*, 2016; Karley *et al.*, 2017), sieved to 3 cm to remove rocks and large debris, and homogenized. The high diversity inoculum consisted of whole soil, and half of the high diversity inoculum was steam sterilized by autoclaving at 121 °C for 3 h, allowing it to cool for 24 h and then autoclaving it again at 121 °C for a further 3 h. The low diversity inoculum was prepared by blending 50 ml of high diversity inoculum with twice the volume of water, filtering the solution through a 38 μm sieve, and vacuum filtering the collected solution through a Whatman filter paper no. 1 (Bennett *et al.*, 2011). Half of the filtrate was autoclaved at 121°C for 20 min. When preparing the experimental pots, those assigned to the high-diversity treatment were inoculated with high-diversity inoculum (whole soil) and autoclaved low-diversity inoculum (filtrate), while those assigned to the low-diversity treatment were inoculated with low-diversity inoculum and autoclaved high-diversity inoculum. This process allowed us to homogenize the abiotic inputs across our pots while varying the microbial community. For further details please see the section below and Figure S2 (Supplementary material).

The filtration process eliminated larger soil microbes, such as arbuscular mycorrhizal (AM) fungi, from the low diversity inoculum. We found significant differences in microbial phylogenetic diversity (see below) between high and low diversity inoculum (Faith’s phylogenetic diversity index 38.31±3.66 for high diversity soil, 8.76±5.94 for low diversity soil; F_1, 10_ = 22.05, *P* < 0.001) which also translated into different microbial diversities in the rhizosphere (see Results below).

### Microcosm setup

Experimental pots (1 L) were assembled as depicted in Fig. S2 (Supplementary material). We added 100 ml of sterile background soil to the bottom and top of each pot to reduce the risk of microbial contamination between pots when watering. Sterile background soil was prepared by mixing Sterilized Loam (Keith Singleton, Cumbria, UK) and sand (ratio 1:1), autoclaving this mixture at 121 °C for 3 h, allowing it to cool for 24 h and then autoclaving it again at 121 °C for a further 3 h. Between the layers of sterile background soil, we added a mix of 100 ml live or sterile high diversity inoculum (10% of the pot volume) and 700 ml of sterile background soil. Pots assigned to the high diversity treatment were filled with live high diversity inoculum and received 1 ml of sterile low diversity inoculum, while pots assigned to the low diversity treatment were filled with sterile high diversity inoculum and received 1 ml of live low diversity inoculum. In this way, we controlled for physical and chemical differences between pots, which only differed in terms of their microbial community. One potato seedling was transplanted into each pot, pots were randomized into two blocks, and left to grow in an insect-screened greenhouse with an average temperature of 25 °C and 16:8 h (light:dark) photoperiod.

Five weeks after transplanting, two apterous adult aphids of *M. euphorbiae* clone AK13/08 were added to 20 plants, two apterous adult aphids of clone AK13/18 were added to 20 plants, and 20 plants were left uninfested. All plants were screened with a microperforated plastic bag (Sealed Air, UK) that allowed transpiration while preventing aphid escape. Three weeks following infestation, we collected from each pot five aphids, leaves, roots and rhizosphere soil (≈500 mg each) and stored them at −80 °C. The five aphids were randomly collected from five different leaves on each plant. Leaf samples were also randomly collected from each plant, being careful to not sample from leaves infested by aphids. Rhizosphere soil was sampled from soil still adhering to the roots after removing loose soil, released by vigorously shaking roots and then sampled. Roots were carefully washed with tap water before collecting a random sample of root tissue. Aphid infestation was scored using a 0–5 scale of severity (scale 0 = no aphids, scale 1 = between 1 and 250 aphids, scale 2 = between 251 and 500 aphids, scale 3 = between 501 and 750 aphids, scale 4 = between 751 and 1,000 aphids, scale 5 = more than 1,000 aphids).

### DNA extraction, Illumina Miseq libraries preparation and sequencing

Samples were crushed in an extraction buffer (10 mM Tris, 100 mM NaCl, 10 mM EDTA, 0.5% SDS) using three 1 mm ∅ stainless steel beads per tube, with the aid of a bead mill homogenizer set at 30 Hz for 5 min (TissueLyzer II, Qiagen, UK). Total DNA was extracted using phenol/chloroform, and it was subsequently checked for quantity and quality with a Nanodrop 2000 (Thermo Fisher Scientific Inc., USA). We conducted a metabarcoding analysis for both bacterial and fungal communities of leaves, roots and rhizosphere soil, and bacterial communities of aphids. Bacterial communities were characterized by targeting the 16S rRNA gene with primers 515f/806rB (Caporaso *et al.*, 2012). Fungal communities were analysed by amplifying the fungal ITS2 region of the rRNA with primers ITS3-KYO/ITS4 (Toju *et al.*, 2012). Amplifications were also carried out on DNA extracted from soil inoculum, and non-template controls where the sample was replaced with nuclease-free water in order to account for possible contamination of instruments, reagents and consumables used for DNA extraction (see Supplementary material).

PCR reactions were performed in a total volume of 25 μl, containing about 50 ng of DNA, 0.5 μM of each primer, 1X KAPA HiFi HotStart ReadyMix (KAPA Biosystems, USA) and nuclease-free water. Amplifications were performed in a Mastercycler Ep Gradient S (Eppendorf, Germany) set at 95 °C for 3 minutes, 98 °C for 30 s, 55 °C for 30 s and 72 °C for 30 s, repeated 35 times, and ended with 10 minutes of extension at 72 °C. Reactions were carried out in technical triplicate, in order to reduce the stochastic variability during amplification (Schmidt *et al.*, 2013), and a no-template control in which nuclease-free water replaced target DNA was utilized in all PCR reactions (see Supplementary material). We also PCR-tested all root samples for the presence of AM fungi using specific primers (Lee *et al.*, 2008), finding presence of AM fungi just in plants grown on high-diversity treatment soil.

Libraries were checked on agarose gel for successful amplification and purified with Agencourt AMPure XP kit (Beckman and Coulter, CA, USA) using the supplier’s instructions. A second short-run PCR was performed in order to ligate the Illumina i7 and i5 barcodes and adaptors following the supplier’s protocol, and amplicons were purified again with Agencourt AMPure XP kit. Libraries were then quantified through Qubit spectrophotometer (Thermo Fisher Scientific Inc., USA), normalized using nuclease-free water, pooled together and sequenced on an Illumina MiSeq platform using the MiSeq Reagent Kit v3 300PE chemistry following the supplier’s protocol.

### Raw reads processing

De-multiplexed forward and reverse reads were merged using PEAR 0.9.1 algorithm using default parameters (Zhang *et al.*, 2014). Data handling was carried out using QIIME 1.9 (Caporaso *et al.*, 2012), quality-filtering reads using default parameters, binning OTUs with a 97% cut-off and discarding chimeric sequences using VSEARCH (Rognes *et al.*, 2016). Singletons and OTUs coming from amplification of chloroplast DNA were discarded from the downstream analyses. Within the ITS2 dataset, all non-fungal OTUs were discarded using ITSx (Bengtsson-Palme *et al.*, 2013). Taxonomy was assigned to each OTU through the BLAST method by querying the SILVA database (v. 132) for 16S (Quast *et al.*, 2012), and UNITE database (v. 8.0) for ITS2 (Nilsson *et al.*, 2019).

### Data analysis

Data analysis was performed using R statistical software 3.5 (R Core Team, 2013) with the packages *phyloseq* (McMurdie and Holmes, 2013), *vegan* (Dixon, 2003) and *picante* (Kembel *et al.*, 2010).

#### Core microbiota

The core microbiota was identified separately for each compartment using the package *ampvis2* (Andersen *et al.*, 2018), considering an OTU as a member of the core microbiota if it was retrieved at a relative abundance of >0.1% in more than 50% of samples. We also tested if the number of core OTUs shared by pairs of compartments was greater/less than an overlap generated by random chance. To do so, for each compartment we draw a group of random OTUs from all those identified in that compartment, in a quantity equal to the number of observed OTUs for that compartment (i.e., if for a compartment we observed 10 core OTUs, we drew 10 random OTUs from the pool of OTUs identified in that compartment). Then, we calculated the number of overlapping random OTUs between all pairs of compartments. We did this separately for bacteria and fungi, and we repeated it 10,000 times. Then, for each pair of compartments, we selected the highest number of random overlapping OTUs among the 10,000 permutations and we tested it against the number of observed overlapping OTUs for that pair of compartments using a χ^2^ test.

#### Phylogenetic diversity

We selected Faith’s phylogenetic diversity index (Faith, 1992) to estimate the diversity of microbiotas in our system because, in contrast to other indices, it takes into account the phylogenetic relationship between taxa within the community. Comparison of diversity indices among groups was performed by fitting a linear mixed-effects model, separately for bacterial and fungal community, specifying *compartment* (i.e., rhizosphere soil, root, leaf, aphid), *soil treatment, plant species, herbivory* (and their interactions) as fixed factors and *aphid clonal line* and *block* as a random effect (Table 1). We also ran separate analyses for plant (rhizosphere soil, root, leaf) and aphids obtaining comparable results (Tab. S1 and S2). The use of *aphid clonal line* as a random variable in the mixed-effects model allowed for the control of differences in the performance of aphid clonal lines. Models were fitted using the *lmer*() function under the *lme4* package (Bates *et al.*, 2015) and the package *emmeans* was used to infer pairwise contrasts (corrected using False Discovery Rate, FDR).

**Table 1.** Models testing the effect of compartment (aphids, leaves, roots, rhizosphere soil), soil treatment (high diversity, low diversity), plant species (*S. tuberosum, S. vernei*), herbivory (infested, control) and their interaction on the phylogenetic diversity (linear mixed-effect model) and taxonomical structure (PERMANOVA) of bacterial and fungal communities.

|  | Bacterial community |  |  |  |  | Fungal community |  |  |  |  |
| --- | --- | --- | --- | --- | --- | --- | --- | --- | --- | --- |
|  |  | Phylogenetic diversity |  | PERMANOVA |  |  | Phylogenetic diversity |  | PERMANOVA |  |
| Factors | df | $\chi^2$ | P | F | P | df | $\chi^2$ | P | F | P |
| Compartment (Cp) | 3 | 843.62 | <0.001 | 23.39 | 0.001 | 2 | 895.91 | <0.001 | 12.9894 | 0.001 |
| Soil treatment (S) | 1 | 2.19 | 0.13 | 13.26 | 0.001 | 1 | 225.62 | <0.001 | 14.906 | 0.001 |
| Plant species (P) | 1 | 18.11 | <0.001 | 2.37 | 0.001 | 1 | 10.67 | <0.01 | 1.9625 | 0.003 |
| Herbivory (H) | 1 | 34.66 | <0.001 | 2.52 | 0.001 | 1 | 47.21 | <0.001 | 3.7912 | 0.001 |
| Cp × S | 3 | 37.23 | <0.001 | 5.67 | 0.001 | 2 | 252.44 | <0.001 | 5.7238 | 0.001 |
| Cp × P | 3 | 8.18 | 0.04 | 1.51 | 0.005 | 2 | 1.66 | 0.43 | 1.3138 | 0.03 |
| S × P | 1 | 23.6 | <0.001 | 1.69 | 0.025 | 1 | 3.74 | 0.05 | 1.7088 | 0.014 |
| Cp × H | 2 | 16.51 | <0.001 | 1.61 | 0.007 | 2 | 20.7 | <0.001 | 2.0339 | 0.001 |
| S × H | 1 | 0.01 | 0.9 | 1.25 | 0.11 | 1 | 5.61 | 0.01 | 1.2592 | 0.107 |
| P × H | 1 | 2.26 | 0.13 | 1.17 | 0.18 | 1 | 0.16 | 0.68 | 1.1442 | 0.217 |
| Cp × S × P | 3 | 12.57 | <0.01 | 1.42 | 0.005 | 2 | 0.28 | 0.86 | 1.34 | 0.029 |
| Cp × S × H | 2 | 6.86 | 0.03 | 1.28 | 0.07 | 2 | 12.38 | <0.01 | 1.107 | 0.211 |
| Cp × P × H | 2 | 1.9 | 0.38 | 1.04 | 0.31 | 2 | 1.18 | 0.55 | 1.0136 | 0.402 |
| S × P × H | 1 | 2.59 | 0.1 | 1.13 | 0.20 | 1 | 1.33 | 0.24 | 1.1739 | 0.152 |
| Cp × S × P × H | 2 | 0.25 | 0.87 | 1.05 | 0.29 | 2 | 0.84 | 0.65 | 1.1249 | 0.186 |

#### Community structure

We analyzed the effects of treatment factors (*compartment, soil treatment, plant species, herbivory* and their interactions) on the structure of the microbial communities using a multivariate approach. Distances between pairs of samples, in terms of community composition, were calculated using a unweighted Unifrac matrix, and then visualized using a Canonical Analysis of Principal Coordinates (CAP) procedure (Anderson and Willis, 2003). Differences between sample groups were inferred through PERMANOVA multivariate analysis (999 permutations stratified at the level of *block* and *aphid clonal line*). The use of *aphid clonal line* for stratification in PERMANOVA allowed for the control of differences in the performance of aphid clonal lines. We also ran separate analyses for plant (rhizosphere soil, root, leaf) and aphids obtaining comparable results (Tab. S1 and S2).

#### Soil diversity vs. plant species vs. herbivory driven effects. Which is strongest?

We assessed the impact of soil treatment, plant species and herbivory for each OTU using the R package *DESeq2* (Love *et al.*, 2014). Using this package, we calculated the effect of each factor (herbivory, plant species, and soil microbial diversity) to the abundance of OTU (expressed as abs log_2_ Fold Changes) in each plant compartment. We first built a model using *compartment* (leaves, roots and rhizosphere), *soil treatment, herbivory* and *plant species* as factors. Then, we extracted the appropriate contrasts (*Low diversity/High diversity* for soil treatment, *S. vernei/S. tuberosum* for plant species and *Herbivore/No herbivore* for herbivore treatment) for each compartment (leaves, roots and rhizosphere). From each contrast, we used the absolute log2 Fold Change values (*ashr* shrinked (Stephens, 2017)) for each OTU to quantify the impact of soil, plant and herbivore treatments on the microbiota in each compartment. Comparisons of absolute log2 Fold Change values were performed by fitting a linear mixed-effects model, specifying *compartment, treatment* (herbivory, plant or soil) and their interaction as fixed factors and *OTU identity* as a random effect, and using the package *emmeans* to infer contrasts (FDR corrected).

#### Aphid infestation

We tested whether the aphid infestation levels were influenced by soil microbial diversity by fitting a cumulative link mixed model using the *ordinal* R package (Christensen, 2015), specifying *soil treatment, plant species*, and their interaction as fixed factors and *block* and *aphid clonal line* as a random effect.

## Results

### Dataset summary and community composition

Overall, we identified 43,879 bacterial and 4,713 fungal OTUs. The analysis of the core microbiota resulted in identifying 150 core bacterial OTUs and 26 fungal OTUs (Fig. 1). We identified 81 bacterial and 19 fungal OTUs as the core microbiota of rhizosphere soil, where the bacterial community was largely dominated by uncultured taxa (50.6%), *Ramlibacter* (10.1%), *Pseudomonas* (9.1%), *Massilia* (8.6%) and *Chitinophaga* (6.1%), while the fungal community was mostly represented by *Peziza* (44.9%), unidentified fungi (18.1%), *Humicola* (9.1%), *Mortierella* (8.8%), *Penicillium* (8.6%), *Mucor* (5.5%) and *Trichoderma* (4.9%). The core microbiota of roots was represented by 74 bacterial and 16 fungal OTUs. Root tissues were mainly associated with *Flavobacterium* (39.4%) and uncultured bacterial taxa (23.7%), while *Peziza* (22.9%), uncultured taxa (18.1%) and *Fusarium* (11.1%) dominated the fungal community. Leaf core microbiota was represented by 20 bacterial and 6 fungal OTUs, with a higher abundance of *Stenotrophomonas* (24.1%), *Candidatus Hamiltonella* (20.9%), *Flavobacterium* (10.5%) and *Pedobacter* (9.4%), while fungi were mainly represented by *Cladosporium* (42.3%), *Penicillum* (34.3%) and *Peziza* (13.6%). Aphids were mainly associated with Buchnera (64.1%) and Candidatus *Hamiltonella* (35.8%). We identified 5 bacterial OTUs (*Flavobacterium* – 2 OTUs, *Citrobacter, Pedobacter, Terrimonas*) shared between all compartments, and 4 fungal OTUs (*Cladosporium, Humicola, Penicillium* and *Peziza*) shared between leaves, roots and rhizosphere. For a more detailed analysis of the core microbiota, please see the Supplementary material.

**Figure 1.**
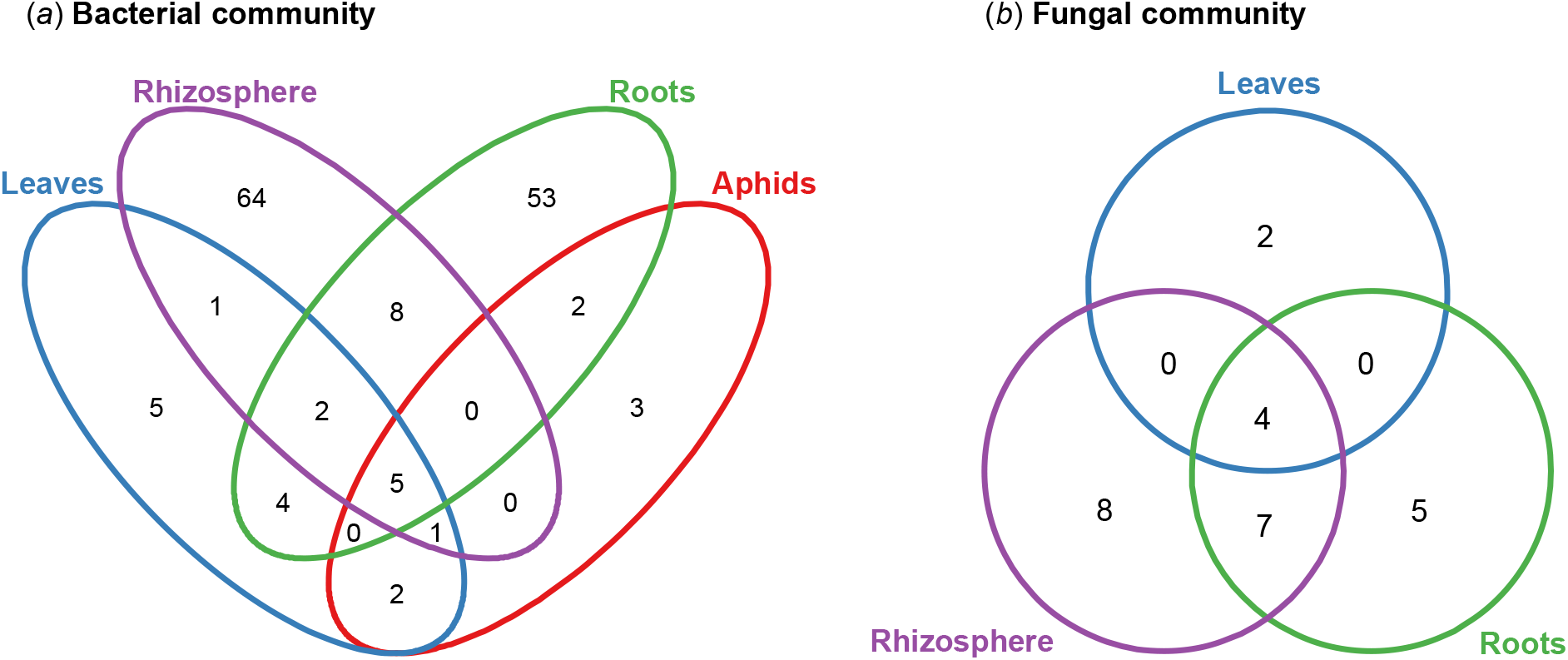
Core bacterial (A) and fungal (B) OTUs shared between compartments. Core OTUs were identified among those with relative abundance of >0.1% in more than 50% of samples.

To gain understanding on the overlap of core OTUs between compartments, we tested the number of observed core OTUs overlapping between two compartments against a the number of OTUs overlapping between two compartments due to random sampling. We found that the overlap of bacterial core OTUs between pairs of plant compartments was always greater than random chance (*P* < 0.05). Aphids also showed a higher number of shared bacterial core OTUs with leaves than random chance (8 OTUs, χ^2^ = 5.4; *P* = 0.01), but not with roots (7 OTUs, χ^2^ = 2.7; *P* = 0.09) and rhizosphere soil (5 OTUs, χ^2^ = 2; *P* = 0.15). In fungi, we saw that the OTUs shared between leaves and roots (χ^2^ = 1.3; *P* = 0.25) or rhizosphere soil (χ^2^ = 0.5; *P* = 0.47) were not different than those shared by chance, while OTUs shared between roots and rhizosphere soil were greater than those shared by chance (χ^2^ = 6.2; *P* = 0.01).

### Phylogenetic diversity

For bacterial communities, we found a significant compartment × soil treatment × plant species interaction (Tab. 1). In all plant compartments (leaves, roots, rhizosphere) we found a higher phylogenetic diversity in *S. vernei* than in *S. tuberosum* when plants were grown on low-diversity soil treatment, and no differences between the two plant species were found when plants were grown on high-diversity soil treatment (Supplementary material Tab. S3). Plant species did not influence aphid bacterial diversity when they were exposed to plants grown on high- or low-diversity soil treatment (Supplementary material Tab. S3). In fungal communities we found a significant effect of the factor “plant species”, reporting a higher diversity in *S. vernei* than *S. tuberosum* plants (*P* = 0.001, Tab. 1), although we did not find any significant interaction with other factors.

We found a significant compartment × soil treatment × herbivory interaction in both bacterial and fungal communities. Post-hoc contrasts show a higher leaf bacterial diversity in aphid-infested plants compared with uninfested control plants when they were grown on low-diversity soil treatment (Supplementary material, Tab. S4). Root bacterial and fungal communities, in both soil treatments, had higher diversity values in infested plants compared with uninfested control plants (Supplementary material, Tab. S4). In the rhizosphere we observed differences between infested and uninfested plants in both bacterial and fungal community diversity of plants grown on high-diversity soil treatment, while this difference was found just in the fungal community of plants grown on low-diversity soil treatment (Supplementary material, Tab. S4).

We found phylogenetic diversity of the aphid microbiota was highest in the low diversity treatment (Figure S3), which mirrored differences in aphid infestation levels (χ^2^=8.19, df=1, *P*=0.004; mean infestation scores: 3.10±0.23 for high diversity soil, 2.45±0.28 for low diversity soil).

### Microbial community composition

The multivariate analysis (i.e., the PERMANOVA) reported a significant compartment × soil treatment × plant species interaction (Tab. 1). Post-hoc contrasts showed differences between *S. vernei* and *S. tuberosum* in the structure of leaf, root and rhizosphere bacterial and fungal communities when plants were grown on low-diversity soil treatment (Fig. 2a-f, Tab. S5). In high-diversity soil treatment, only root fungal communities differed between *S. vernei* and *S. tuberosum* (Fig. 2e, Supplementary material Tab. S5). On the other hand, differences between low-diversity and high-diversity treatments were found in both *S. vernei* and *S. tuberosum* in root and rhizosphere communities (both bacterial and fungal, Fig. 2b-c and 2e-f, Supplementary material Tab. S6). In leaves, differences between soil treatments were found just in the bacterial community of *S. vernei* (Fig. 2a, Supplementary material Tab. S6). We also found a significant compartment × herbivory interaction (Tab. 1), with herbivory influencing bacterial communities in all compartments, but fungal community just in leaves and roots (Tab. 2). Our multivariate analysis demonstrated that the strongest driver of rhizosphere, root and leaf microbial community structure was soil diversity treatment (Tab. 2) for both bacterial and fungal communities. Indeed, bacterial and fungal communities responded to soil diversity treatment and plant species across all compartments (roots, rhizosphere and leaves) (Table 2, Fig. 2a-f). Based on the variation explained by each factor included in the model, soil microbial diversity was the most important factor shaping the microcosm’s microbiota in all compartments (Table 2). The variation explained by soil microbial diversity tended to decrease when moving across compartments from rhizosphere to leaves and aphids (Table 2).

**Table 2.**
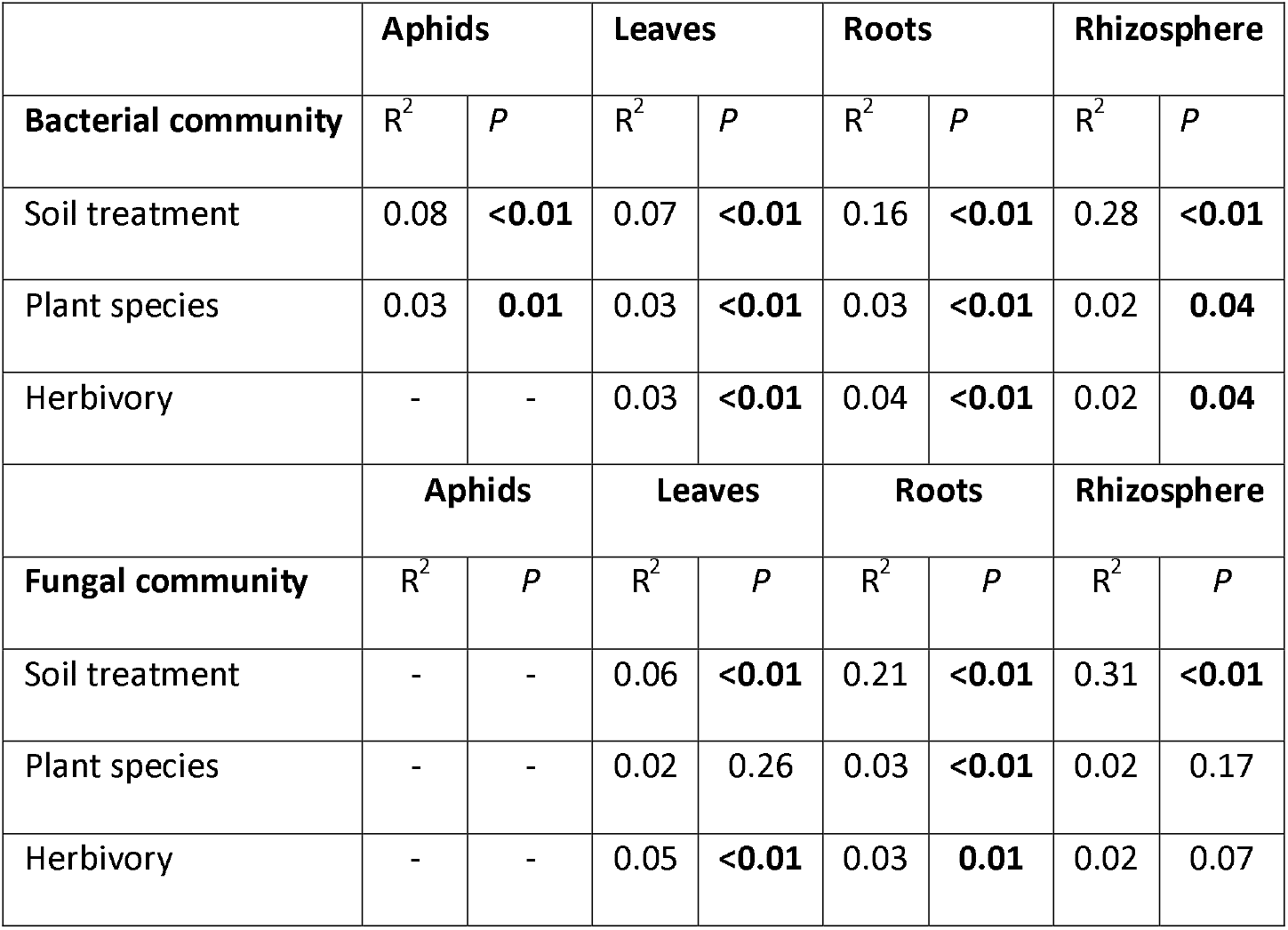
Analysis of the effects of soil treatment (high diversity, low diversity), plant species (*S. tuberosum, S. vernei*), herbivory (infested, control) on the bacterial and fungal community taxonomical structure for each compartment (aphids, leaves, roots and rhizosphere) performed through PERMANOVA (Unifrac distance matrix). Analyses were run separately for each compartment within each community. Values in bold represent *P*<0.05.

|  | Aphids |  | Leaves |  | Roots |  | Rhizosphere |  |
| --- | --- | --- | --- | --- | --- | --- | --- | --- |
| **Bacterial community** | R<sup>2</sup> | P | R<sup>2</sup> | P | R<sup>2</sup> | P | R<sup>2</sup> | P |
| Soil treatment | 0.08 | **<0.01** | 0.07 | **<0.01** | 0.16 | **<0.01** | 0.28 | **<0.01** |
| Plant species | 0.03 | **0.01** | 0.03 | **<0.01** | 0.03 | **<0.01** | 0.02 | **0.04** |
| Herbivory | - | - | 0.03 | **<0.01** | 0.04 | **<0.01** | 0.02 | **0.04** |
|  | Aphids |  | Leaves |  | Roots |  | Rhizosphere |  |
| **Fungal community** | R<sup>2</sup> | P | R<sup>2</sup> | P | R<sup>2</sup> | P | R<sup>2</sup> | P |
| Soil treatment | - | - | 0.06 | **<0.01** | 0.21 | **<0.01** | 0.31 | **<0.01** |
| Plant species | - | - | 0.02 | 0.26 | 0.03 | **<0.01** | 0.02 | 0.17 |
| Herbivory | - | - | 0.05 | **<0.01** | 0.03 | **0.01** | 0.02 | 0.07 |

**Figure 2.**
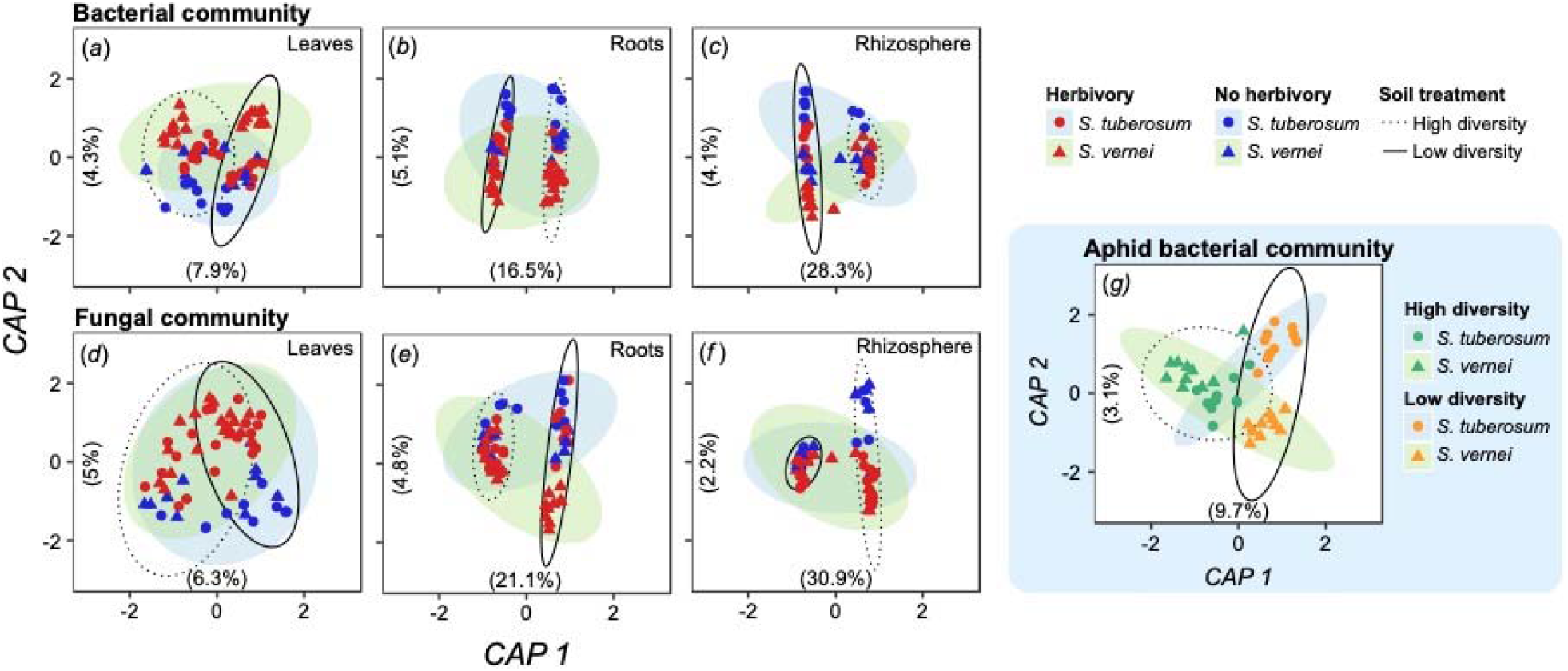
Canonical Analysis of Principal coordinates (CAP) analysis of bacterial and fungal communities Unifrac distance matrix for each compartment. We report the response of bacterial (*a-c*) and fungal (*d-f*) communities to soil microbial diversity, plant species and herbivory in leaves (*a, d*), roots (*b, e*) and rhizosphere soil (*c, f*). Aphid bacterial community (*g*) responded to both plant species and soil microbial diversity. For each graph, percentages in parentheses inside each graph along the axes report the variance explained by the respective axis.

### Soil diversity vs. plant species vs. herbivory driven effects. Which is strongest?

We answered this question in two ways, focusing on the single factors included in our design (soil treatment, plant species, herbivory). First, as discussed above, using the variation explained by each of our predictor variables in our PERMANOVA model, we determined that the predictor that explained the most variation was the initial soil community diversity. Both soil diversity, plant species and herbivory influenced bacterial and fungal assemblies in our system. Soil treatment explained ≈30% (rhizosphere), ≈20% (root), and ≈7% (leaf) of variation in microbiome community composition (Table 2). However, the variance in community composition explained by both plant species and herbivory (3–5%) was always lower than the variance explained by the soil treatment. Furthermore, soil treatment explained ≈8% of variation in aphid microbiota (Table 2). This suggests that the soil-driven effect is stronger than the other effects in our system.

Second, to investigate in more detail which factor (soil, plant, herbivore) had a stronger influence on plant microbiome composition we examined the magnitude of change in abundance for each OTU (absolute log_2_ Fold Changes) in relation to soil treatment, plant species and herbivory. For both bacterial and fungal communities, and in all compartments, the changes produced by soil treatment were greater than those produced by herbivory and plant species (χ^2^_bacteria_ =23331.3 and χ^2^_fungi_ =1055, df=2, P<0.001, Fig. 3), with the only exception being the leaf fungal community, where no differences were found between the three factors (Fig. 3b). Also, in all cases, there was no difference between the changes produced by herbivory and those produced by plant species (Fig. 3). The analysis of changes in the abundance of OTUs in aphids revealed that soil diversity treatment had a greater influence than plant species in shaping aphid bacterial communities (χ^2^=766.8, df=1, *P*<0.001; abs(log_2_FoldChange)_soil_ = 1.7±0.1 and abs(log_2_FoldChange)_plant_ = 0.37±0.1). Collectively, these results demonstrate that the strongest effect on microbial taxa in the rhizosphere, roots, shoots, and aphid herbivores is driven by the initial soil community diversity.

**Figure 3.**
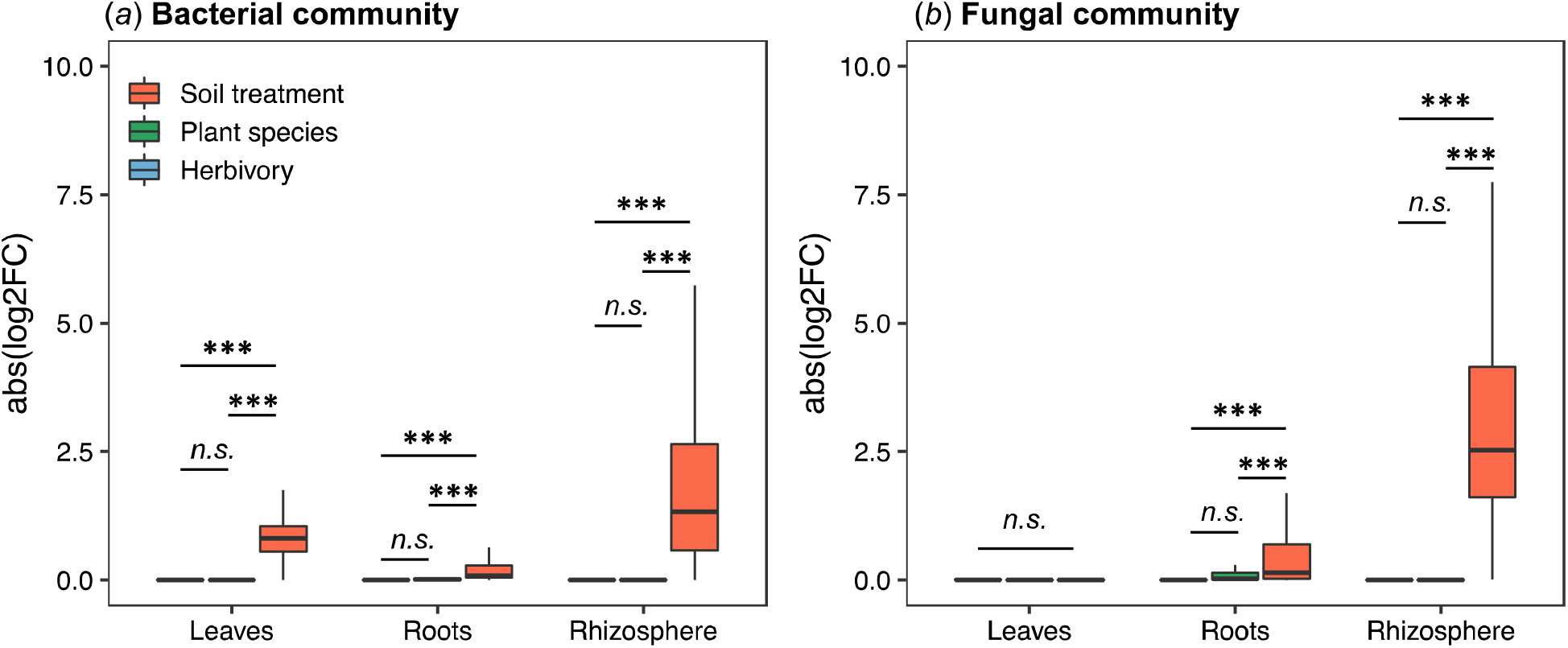
Magnitude of changes in abundance for each OTU (absolute log_2_ Fold Changes). For each compartment (leaves, roots and rhizosphere) we investigated the response of single OTUs to soil microbial diversity (red), plant species (green) and herbivory (blue), for (*a*) bacterial and (*b*) fungal communities. Comparisons were tested using a linear mixed-effect model, and contrasts were extracted using the function emmeans. *** *P*<0.001.

## Discussion

Here we test the influence of multiple drivers (and their interaction) on plant microbiome diversity and composition, and we show that, of all the drivers tested, soil microbial diversity had the greatest influence on the microbial community composition of rhizosphere, roots, leaves, and even aphid herbivores. Thus, we correctly hypothesized that soil microbial diversity drives changes in plant and herbivore microbiota and that this effect would be much stronger than plant species or herbivory. This influence of soil microbial diversity correlated with aphid abundance on infested plants. Furthermore, we showed herbivory and plant species also affect the microbiome community composition of leaves, roots and rhizosphere, but their effects are weaker than those driven by soil diversity. We also observed that the response of plant microbiome to herbivory or plant species differs according to soil treatment.

Plant and insect microbial diversity was influenced by soil microbiota and interactions between soil microbiota and plant species and herbivory. When we quantified the relative contribution of the main effects (soil, plant species, and herbivory), soil treatment was the strongest driver of plant related microbiota composition. Soil community composition is well-known to influence plant microbial composition (Bulgarelli *et al.*, 2012; Schlaeppi *et al.*, 2014; Bai *et al.*, 2015; Zarraonaindia *et al.*, 2015; de Souza *et al.*, 2016; Wagner *et al.*, 2016; Mezzasalma *et al.*, 2018; Grady *et al.*, 2019), but here we reveal that soil produces a stronger effect when compared to other factors (i.e. plant species and herbivory). Soil represents a major reservoir of microbial inoculum for plants (Zarraonaindia *et al.*, 2015; Trivedi *et al.*, 2020), especially belowground. However, we also observed that the effect driven by soil is a function of plant species, likely because the intrinsic characteristics of each plant species can modulate specific changes in the diversity and composition of microbial communities at each compartment (Turner *et al.*, 2013; Dastogeer *et al.*, 2020).

Interestingly, we found that soil microbial diversity influences the phylogenetic diversity and structure of aphid microbiota. The influence of the soil microbial diversity on aphid bacterial communities and aphid infestation level could potentially be explained through two, non-mutually exclusive, mechanisms: (i) translocation of microbes from the rhizosphere through or on the plant and (ii) changes in plant physiology and/or metabolome. Although leaves are physically separated from roots, their microbiomes can still interact at interfaces such as the stem, and microbial translocation could occur due to active and passive mechanisms (Bai *et al.*, 2015; Wagner *et al.*, 2016). A recent study comparing the microbiota of caterpillars feeding on detached leaves and intact plants found the microbiota of caterpillars that fed on intact plants had a similar community composition to the soil microbiota (Hannula *et al.*, 2019) suggesting direct (splashing of soil microbiota on leaves) or indirect (movement through the plant) microbial translocation. However, our data does not show this pattern, as the core microbiome belowground is different from shoots and herbivores, few OTUs are common to all compartments in the system, and the overlap of core OTUs between aphids and belowground compartments is not greater than random chance. The aphids in our system employ a different feeding strategy (sap-feeding) compared to the caterpillars (chewing) in the previous study (Hannula *et al.*, 2019), and chewing herbivores may have an increased likelihood of environmental uptake of microbes (Paniagua Voirol *et al.*, 2018). We thus find it unlikely that microbes in our system were translocated through or on the plant to the herbivore.

The second potential mechanism is through changes in plant physiology. Many soil microbes are able to modulate plant nutrient intake, or prime plant defences (Martinez-Medina *et al.*, 2016). The composition of belowground microbial communities can alter plant metabolism (Li *et al.*, 2019), which in turn influences herbivore fitness (Mason *et al.*, 2019). Our low diversity treatment lacked large microbes including arbuscular mycorrhizal fungi, a group well known to prime plant defences (Jung *et al.*, 2012; Bennett *et al.*, 2018), although aphids are less susceptible to changes in defenses primed by arbuscular mycorrhizal fungi (Koricheva *et al.*, 2009). Thus, the changes we observed in the aphid microbiome could be due to changes in host plant physiology and metabolome, for example triggering of plant defences which has been shown to decrease the diversity of plant-associated microbial communities (Kniskern *et al.*, 2007). The higher aphid abundance on plants grown in the high diversity soil provides indirect evidence for such changes in plant biochemical composition, as does a previous study showing increased aphid suitability as a host for parasitoids when feeding on Solanum plants grown with AM fungi from the same site (Bennett *et al.*, 2016). While no previous study found a relationship between soil microbial diversity and aphid infestation, previous studies show that soil microbiota can influence insect feeding behaviour through changes in the plant metabolome (Badri *et al.*, 2013), which can represent the likely mechanism driving our observations.

While a clear consumer-driven effect was observed on the plant microbiome in our study, it was a weaker effect than soil microbial diversity. Thus, herbivory plays a less significant role in determining plant microbiome composition. The herbivory-driven effect on the bacterial community composition of the roots and rhizosphere could be driven by changes in plant physiology (e.g. defence activation, carbon metabolism) and root exudation due to aphid feeding activity (Züst and Agrawal, 2016; Hoysted *et al.*, 2018). Herbivory has been shown to alter the types of organic compounds released at the root surface leading to changes in the composition of rhizosphere microbial communities (Lareen *et al.*, 2016; Hu *et al.*, 2018). Previous research has shown that Bemisia tabaci (whitefly) herbivory can alter the rhizosphere microbiome of pepper plants (Kong *et al.*, 2016), and artificial induction of plant defences (Hein *et al.*, 2008), or their deactivation (Kniskern *et al.*, 2007), has been shown to shape rhizosphere microbial communities. In our study we focused on a sap-feeding insect, which is known to trigger specific physiological responses in plants. Thus, further research is needed to test if different herbivores with different feeding strategies are able to drive changes in the microbiota thriving in different plant compartments.

Plant species was also a predictor shaping the microbiome community composition of the rhizosphere, plants, and herbivores in our study. Plant species is known to be a strong driver of root and rhizosphere microbiota because plant species differ in morphology, chemistry and relationship with microorganisms (Trivedi *et al.*, 2020). *Solanum* plants are known to produce toxic glycoalkaloids as chemical defense against herbivory (Altesor *et al.*, 2014), and these compounds could also shape the microbiota associated with the plant. The strength of this effect might be context-dependent due to the interaction with the soil microbial composition (Bulgarelli *et al.*, 2013; Fierer, 2017). This might explain why we observed a greater impact of soil treatment than plant species in the belowground microbiotas in our study. Also, it might explain why we found differences in the bacterial community between plant species just in one soil treatment. It has been previously shown that soil represents a reservoir for leaf microbial communities and that phyllosphere habitat selects for specific members (Grady *et al.*, 2019), which partially explains our observation that the impact of soil microbial diversity was greater than plant species on the phyllosphere bacterial and fungal communities. The differences in the microbiome composition of aphids feeding and reproducing on the two different plant species is not surprising, as it is well known that the identity of the host plant is a major factor in shaping insect-associated microbial community composition (Colman *et al.*, 2012; Malacrinò, 2018). The fact that seeds were not surface-sterilized might be a caveat of our study, potentially influencing the structure of plant microbiome differently for each species. However, previous studies show that even surface-sterilized seeds show differences in their endophytic microbiome according to the plant genotype (Walitang *et al.*, 2018; Raj *et al.*, 2019; Liu *et al.*, 2020), and our results show a major effect driven by soil microbial community composition, thus we are confident that our results were not biased by not surface-sterilizing seeds.

By quantifying and comparing the relative strength of multiple biotic drivers of microbial community composition, our work contributes to a more comprehensive understanding of the factors determining the outcome of plant–microbe–insect interactions, and how plant-associated microbiomes assemble and respond to resource- and consumer-driven effects. Thus, if understood and managed correctly, these interactions have potential to be applied in natural and managed systems to improve food security and safety, or the success of ecological restoration efforts.

## Declarations

### Availability of data and material

Raw data is available at the NCBI SRA database under accession number PRJNA557499.

### Authors’ contributions

All authors designed the experiment. AM and AEB performed the greenhouse experiment. AM prepared library and performed data analysis. AM wrote the first draft of the manuscript and all authors contributed substantially to revisions.

## Acknowledgements

We would also like to thank Davide Bulgarelli (University of Dundee, UK), Marta Jarzyna (Ohio State University, USA), Jeff Powell (Hawksbury Institute for the Environment, Western Sydney University, AU), and Philip Smith (Dundee, UK) for helpful comments on the manuscript.

## Supplementary material

**Supplementary material.** Supplementary results, tables and figures.

