## Supplementary Material for "Soil microbial diversity impacts plant microbiota more than herbivory"

### Supplementary material
#### Soil microbial diversity impacts plant and herbivore microbiomes more than herbivory

Antonino Malacrinò , Alison J. Karley , Leonardo Schena , and Alison E. Bennett

*Journal XXXX, Year YYYY, DOI ZZZ*

### 1 Supplementary results

#### 1.1 Soil inoculum and negative controls

As part of the workflow we included two additional groups of samples for sequencing: soil inoculum and negative control samples. The first group informed us on the microbial community that was inoculated in our microcosm system. The negative control group helped us to account for potential contaminations.

##### 1.1.1 Inoculum

In order to provide a more detailed information of the diversity of our inoculum, we performed the entire molecular workflow on the soil used as microbial inoculum with the same protocol used for experimental samples. This soil was collected during the experimental setup and stored at -80°C until processed.

The bacterial community of soil inoculum was mainly characterized by unidentified Acidobacteria (33.72±0.03%) followed by 19 lineages representing about 41% of the bacterial diversity, while the fungal community was dominated by unidentified fungi (50±5%), *Penicillium* (15±3.7%), *Sclerotinia* (15±15%), uncultured *Schizothecium* (10±10%), *Mucor* (5±5%) and uncultured *Peziza* (5±5%). We observed an overlap between the inoculum and the rhizosphere soil collected at the end of the experiment, for both high-diversity and low-diversity treatments and for both bacterial and fungal community (Fig. S1). We also observed distinct communities for both inoculum and rhizosphere soil (Fig. S1), which is expected as consequence of the evolution of microbial communities during the experiment and the input of microbes from the environment (e.g., seedlings, greenhouse, manipulation).

The community of AM fungi was characterized using a specific set of primers, targeting the 18S region since they are difficult to be amplified using ITS region (Van Geel *et al.*, 2014). In this case, we sequenced both soil inoculum and spore extracted using the methodology suggested by Daniels and Skipper (1982), in order to have a higher resolution. The inoculum community was represented by *Claroideoglomus* (37.84±0.5%), *Glomus* (24.21±0.4%), *Archaeospora* (25.94±4.4%), *Paraglomus* (10.82±1.19%) and *Diversispora* (1.19±0.6%). An additional analysis on spores extracted from soil led to the identification of further lineages of AM fungi: *Gigaspora*, *Scutellospora*, *Ambispora* and unidentified *Glomeromycota.*

##### 1.1.2 Negative control samples

Specifically, we included two categories of control samples:

2. **Non-template control (NTC)**. We included a non-template control (molecular biology grade water instead of DNA sample) for each PCR plate, which was amplified in technical triplicates as carried out for all biological samples. If amplification of NTC was observed, the entire plate was discarded. It is important to note that the PCR master mix used in each well of each plate came from the same pool.

Although no amplification was observed in any sample belonging to both categories, PCR products were processed and sequenced anyway. For most of samples we did not retrieve any sequence, while for a few of them (6 for 16S, 7 for ITS), we retrieved a maximum of 12 sequences per samples that did not pass QC or singletons removal. Therefore, these sequences are likely to come from index hopping, which is known to occur in Illumina machines. We can, therefore, conclude that no contamination was observed in our experiment.

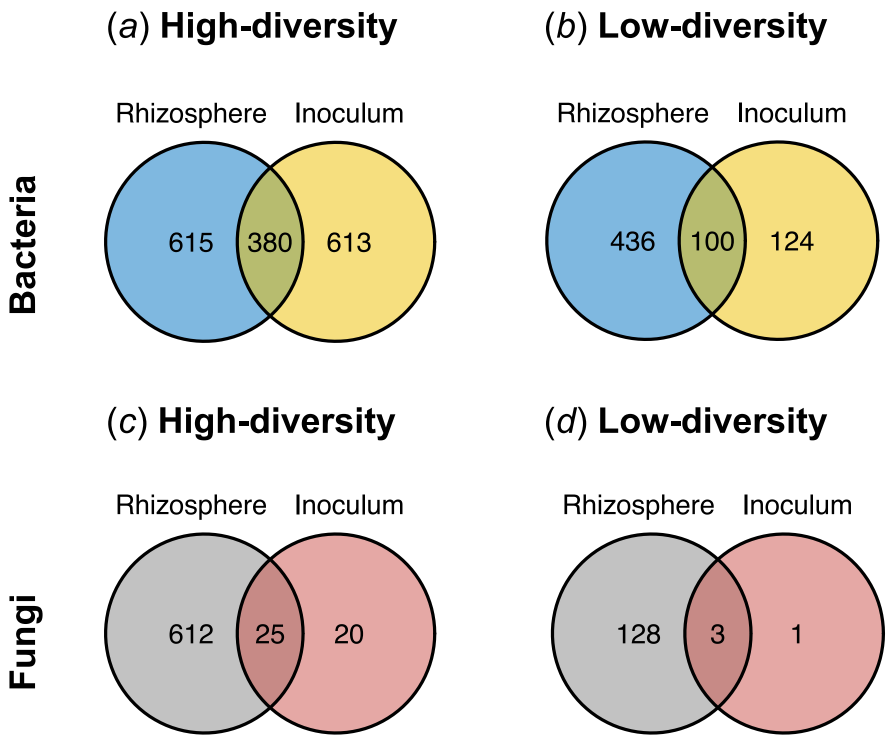


**Figure S1**. Shared bacteria (a, b) and fungal (c, d) OTUs between the rhizosphere soil collected at the end of the experiment for high-diversity (a, c) and low-diversity (b, d) treatments.

##### 1.2 Core community composition

The core bacterial community of rhizosphere accounted for 81 OTUs (Fig. 4a), mainly represented by Proteobacteria (47 OTUs) followed by Verrucomicrobia (7 OTUs), Acidobacteria (7 OTUs), Bacteroidetes (7 OTUs), Planctomycetes (7 OTUs), Actinobacteria (5 OTUs) and Gemmatimonadetes (1 OTU). Roots, however, were highly abundant in Bacteroidetes (61 OTUs) followed by Actinobacteria (8 OTUs), Proteobacteria (3 OTUs) and Verrucomicrobia (2 OTUs). Rhizosphere and roots shared 15 core OTUs, including Bacteroidetes (2 Flavobacterium, 2 Pedobacter, Terrimonas, Flavitalea and an uncultured Microscillaceae), Actinobacteria (2 Nocardioides, Agromyces, Galbitalea and

Pseudoarthrobacter), Proteobacteria (Arenimonas and Citrobacter) and Verrucomicrobia (Lacunisphaera). On leaves the core community (20 OTUs) was equally shared by Actinobacteria (6 OTUs), Bacteroidetes (7 OTUs) and Proteobacteria (7 OTUs). This compartment shares just a single OTU with soil (*Massilia* sp.). Leaves shared 9 OTUs with the belowground compartment: *Citrobacter*, *Flavobacterium* (2 OTUs), *Massilia*, *Nocardioides*, *Pedobacter*, *Pseudoarthrobacter*, *Pseudomonas* and *Terrimonas*. Aphids harboured 13 OTUs in their core microbiome: *Bacillus*, *Buchnera* (2 OTUs), *Hamiltonella*, *Citrobacter*, *Dyadobacter*, *Flavobacterium* (2 OTUs), *Paenibacillus*, *Pedobacter*, *Pseudoflavitalea*, *Pseudomonas* and *Terrimonas*.

On the fungal side (Fig. 4b), the rhizosphere core community (19 OTUs) was dominated by *Penicillium* (4 OTUs), followed by *Trichoderma* (2 OTUs), *Fusarium* (2 OTUs), *Chaetomium* (1 OTU), *Cladosporium* (1 OTU), *Didymella* (1 OTU), *Humicola* (1 OTU), *Mortierella* (1 OTU), *Peziza* (1 OTU) and 5 unidentified fungi. Root fungal community was also dominated by *Penicillium* (3 OTUs) followed by *Peziza* (2 OTU), *Humicola* (2 OTU), *Chaetomium* (1 OTU), *Cladosporium* (1 OTU), *Fusarium* (1 OTUs), *Mucor* (1 OTU), *Olpidium* (1 OTU), *Trichoderma* (1 OTU), *Rhexocercosporidium* (1 OTU) and 2 unidentified fungi. Of these, *Chaetomium*, *Cladosporium*, *Fusarium*, *Humicola*, *Penicillium*, *Peziza*, *Trichoderma* and 2 unidentified fungi were shared between rhizosphere and roots. Leaf core community was dominated by *Cladosporium* (3 OTUs), *Humicola* (1 OTU), *Penicillium* (1 OTU) and *Peziza* (1 OTU) which, excluding 2 *Cladosporium* OTUs, were all shared across the three compartments.

#### 2 Supplementary tables

**Table S1**. Models testing the effect of plant compartment (leaves, roots, rhizosphere soil), soil treatment (high diversity, low diversity), plant species (S. tuberosum, S. vernei), herbivory (infested, control) and their interaction on the phylogenetic diversity (linear mixed-effect model) and taxonomical structure (PERMANOVA) of plant bacterial communities.

| | | **Phylogenetic diversity** | | **PERMANOVA** | |
|---|---|---|---|---|---|
| Factor | df | $\chi^2$ | $P$ | F | $P$ |
| Compartment (Cp) | 2 | 401.82 | **<0.001** | 30.96 | **<0.001** |
| Soil treatment (S) | 1 | 0.14 | 0.7 | 18.8 | **<0.001** |
| Plant species (P) | 1 | 21.03 | **<0.001** | 2.69 | **<0.001** |
| Herbivory (H) | 1 | 17.94 | **<0.001** | 2.83 | **0.003** |
| Cp × S | 2 | 27.05 | **<0.001** | 7.62 | **<0.001** |
| Cp × P | 2 | 2.72 | 0.25 | 1.59 | **0.012** |
| S × P | 1 | 24.04 | **<0.001** | 1.96 | **0.013** |
| Cp × H | 2 | 13.71 | **0.001** | 1.81 | **0.003** |
| S × H | 1 | 0.01 | 0.91 | 1.34 | 0.11 |
| P × H | 1 | 1.88 | 0.16 | 1.22 | 0.17 |
| Cp × S × P | 2 | 4.53 | 0.1 | 1.44 | **0.03** |
| Cp × S × H | 2 | 5.67 | 0.05 | 1.38 | **0.03** |
| Cp × P × H | 2 | 1.58 | 0.45 | 1.09 | 0.29 |
| S × P × H | 1 | 2.16 | 0.14 | 1.17 | 0.21 |
| Cp × S × P × H | 2 | 0.21 | 0.89 | 1.09 | 0.25 |

**Table S2**. Models testing the effect of soil treatment (high diversity, low diversity), plant species (S. tuberosum, S. vernei) and their interaction on the phylogenetic diversity (linear mixed-effect model) and taxonomical structure (PERMANOVA) of aphid bacterial communities.

| | | **Phylogenetic diversity** | | **PERMANOVA** | |
|---|---|---|---|---|---|
| Factor | df | $\chi^2$ | *P* | F | *P* |
| Soil treatment | 1 | 77.16 | **<0.001** | 3.8 | **<0.001** |
| Plant species | 1 | 5.53 | **0.01** | 1.46 | **0.025** |
| Soil treatment × Plant species | 1 | 0.0005 | 0.98 | 1.34 | **0.042** |

**Table S3**. Specific contrasts (*S. tuberosum – S. vernei*) extracted from the linear mixed-effects model to analyse, for each soil treatment, the effect of plant species on the bacterial diversity for each compartment.

| **Compartment** | **Soil treatment** | **Estimate** | ***P*** |
|---|---|---|---|
| Aphids | Low diversity | 4.04 | 0.10 |
| | High diversity | 3.96 | 0.11 |
| Leaves | Low diversity | -14.80 | **0.03** |
| | High diversity | -5.98 | 0.39 |
| Roots | Low diversity | -34.47 | **<0.001** |
| | High diversity | 7.18 | 0.31 |
| Rhizosphere | Low diversity | -34.70 | **<0.001** |
| | High diversity | -6.62 | 0.34 |

**Table S4**. Specific contrasts (*Control – Infested*) extracted from the linear mixed-effects model to analyse, for each soil treatment, the effect of herbivory on bacterial and fungal diversity for each compartment.

| | | **Bacteria** | | **Fungi** | |
|---|---|---|---|---|---|
| Compartment | Soil treatment | Estimate | *P* | Estimate | *P* |
| Leaves | Low diversity | -16.32 | **0.04** | -2.96 | 0.31 |
| | High diversity | 2.34 | 0.75 | -1.74 | 0.56 |
| Roots | Low diversity | -34.01 | **<0.001** | -6.40 | **0.03** |
| | High diversity | -36.67 | **<0.001** | -7.05 | **0.02** |
| Rhizosphere | Low diversity | -14.93 | 0.05 | -6.73 | **0.02** |
| | High diversity | -32.98 | **<0.001** | -24.20 | **<0.001** |

**Table S5**. Specific contrasts (*S. vernei* / *S. tuberosum*) extracted from the PERMANOVA model to analyse, for each soil treatment, the effect of plant species on the bacterial and fungal community structure for each compartment.

| Compartment | Soil treatment | Bacteria ($P$ value) | Bacteria ($P$ value) |
|---|---|---|---|
| Leaves | Low diversity | **0.04** | **0.02** |
| | High diversity | 0.06 | 0.10 |
| Roots | Low diversity | **0.001** | **0.001** |
| | High diversity | 0.22 | **0.002** |
| Rhizosphere | Low diversity | **0.001** | **0.001** |
| | High diversity | 0.3 | 0.05 |

**Table S6**. Specific contrasts (*Low diversity* / *High diversity*) extracted from the PERMANOVA model to analyse, for each plant species, the effect of soil treatment on the bacterial and fungal community structure for each compartment.

| Compartment | Soil treatment | Bacteria ($P$ value) | Bacteria ($P$ value) |
|---|---|---|---|
| Leaves | *S. tuberosum* | 0.07 | 0.10 |
| | *S. vernei* | **0.01** | 0.29 |
| Roots | *S. tuberosum* | **0.001** | **0.001** |
| | *S. vernei* | **0.001** | **0.001** |
| Rhizosphere | *S. tuberosum* | **0.001** | **0.001** |
| | *S. vernei* | **0.001** | **0.001** |

### 3 Supplementary figures

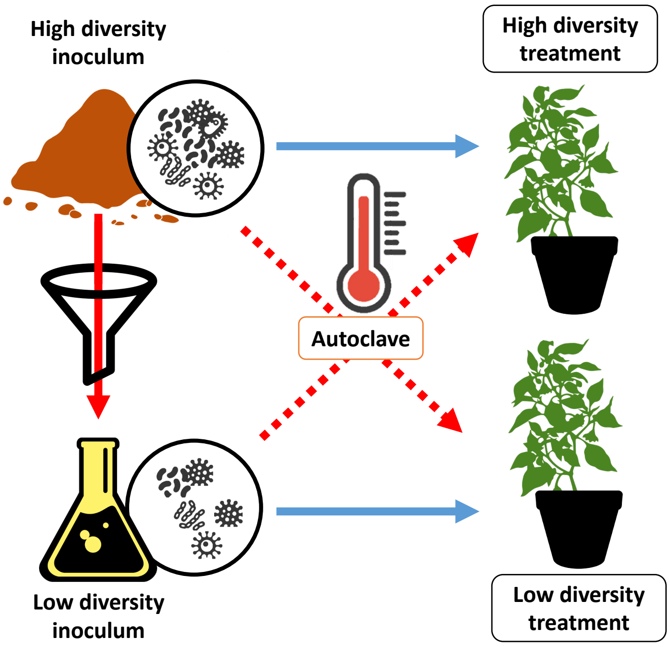


**Figure S2**. Graphical representation of the microcosm setup. High diversity treatment was obtained by mixing the high-diversity inoculum with autoclaved low-diversity inoculum. Low diversity treatment was obtained by mixing the low-diversity inoculum with autoclaved high-diversity inoculum.

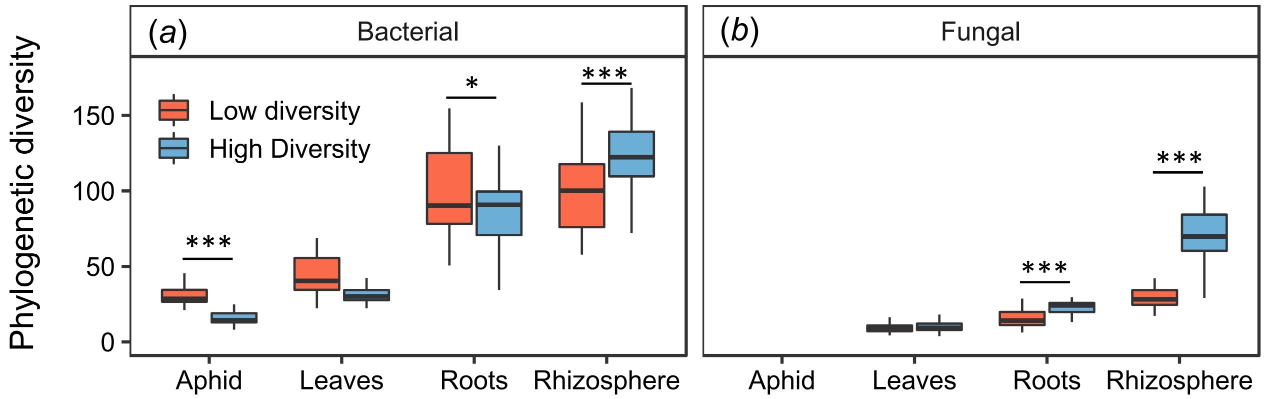


**Figure S3**. Phylogenetic diversity of (a) bacterial and (b) fungal communities in each compartment in response to soil microbial diversity. *** $P<0.001$; * $P<0.05$
